# Response to Problems in interpreting and using GWAS of conditional phenotypes illustrated by alcohol GWAS

**DOI:** 10.1101/290965

**Authors:** Toni-Kim Clarke, Andrew M. McIntosh

## Introduction

In recent correspondence, Holmes and Davey Smith highlight ‘Problems in interpreting and using GWAS of conditional phenotypes illustrated by ‘alcohol GWAS’^1^. The authors suggest that a negative genetic correlation between BMI and alcohol consumption, which we previously reported in the UK Biobank sample, is spurious^2^. In regards to the approach we used to run our genome-wide association study (GWAS) of alcohol consumption adjusted for age and weight, the authors state that the negative genetic correlation with BMI was ‘induced by the very nature of their statistical model’. We agree that their commentary highlights a potential difficulty in epidemiological research. Conditioning a test of association on a collider may introduce spurious correlation. Whilst this may be a general consideration, we suggest that collider bias does not explain the association presented in Clarke et al (2017)^2^.

## Methods

In order to test Holmes and Davey Smith’s assertion, we repeated our GWAS of alcohol consumption in the same 112,117 UK Biobank individuals using the same phenotype, unadjusted for weight. We then took the GWAS summary statistics and, using LD score regression^3^ implemented in LDHub^4^, re-analysed the genetic correlation with BMI.

## Results

Previously, we reported a negative genetic correlation between BMI and alcohol consumption when adjusted for weight (r_g_=−0.15, p=5 × 10^−4^). In our new analysis, the genetic correlation between alcohol consumption and BMI, unadjusted for weight, was still negative but was of even greater magnitude than in the adjusted analysis (r_g_=−0.21, p=7 × 10^−7^). Therefore, the negative genetic correlation between BMI and alcohol consumption becomes attenuated when adjusted for weight, albeit these two estimates are not significantly different from one another (Z=-0.99, p=0.32).

## Discussion

Further independent data also support our assertion that the findings presented in Clarke et al are robust. A negative genetic correlation between BMI and alcohol consumption was reported in a genetic study of AUDIT scores in individuals from the 23andMe cohort^5^. The AUDIT is a 10-item questionnaire designed to measure alcohol consumption and problematic drinking. Conducting a GWAS of AUDIT scores in 20,328 individuals, the authors also report a negative genetic correlation between BMI and AUDIT scores of -0.25 (p=1.5 x 10^−4^), similar in magnitude to our own findings.

The negative genetic correlation between BMI and alcohol consumption, and the positive correlation with education and HDL cholesterol, led us to state in our original publication that alcohol consumption is correlated with ‘many positive health and behavioural traits’. Some of these relationships may be causal; in the case of HDL cholesterol independent studies would suggest that this is in fact the case^6, 7^. However, the positive genetic correlation between alcohol and education, and the negative genetic correlation with BMI, do not lend themselves to straightforward causal interpretation. We accordingly resisted any temptation to draw any causal inferences from these associations.

In summary, in contrast to Holmes and Davey Smith we are less confident that the negative genetic correlation between BMI and consumption is due to conditioning on a collider. Volunteer participation in cohort studies may lead to individuals being atypical of the populations they represent. Further work, from independent data sources and methodologies, is needed to further understand the associations presented in GWAS studies and move towards more robust causal inferences.

